# Are cell wall traits a component of the succulent syndrome?

**DOI:** 10.1101/2022.06.25.497541

**Authors:** Marc Fradera-Soler, Alistair Leverett, Jozef Mravec, Bodil Jørgensen, Anne M. Borland, Olwen M. Grace

## Abstract

Succulence is an adaptation to low water availability characterised by the presence of water-storage tissues that alleviate water stress under low water availability. The succulent syndrome has evolved convergently in over 80 plant families and is associated with anatomical, physiological and biochemical traits. Despite the alleged importance of cell wall traits in drought responses, their significance in the succulent syndrome has long been overlooked. Here, by analysing published pressure–volume curves, we show that elastic adjustment, whereby plants change cell wall elasticity, is uniquely beneficial to succulents for avoiding turgor loss. In addition, we used comprehensive microarray polymer profiling (CoMPP) to assess the biochemical composition of cell walls in leaves. Across phylogenetically diverse species, we uncover several differences in cell wall biochemistry between succulent and non-succulent leaves, pointing to the existence of a ‘succulent glycome’. We also highlight the glycomic diversity among succulent plants, with some glycomic features being restricted to certain succulent lineages. In conclusion, we suggest that cell wall biomechanics and biochemistry should be considered among the characteristic traits that make up the succulent syndrome.

## Introduction

Climate change-induced aridity is expected to increase across much of the globe in the future (Sheffield and Wood, 2008; Jiao et al., 2021). Consequently, it has become imperative that we understand the ways in which plants cope with drought (Choat et al., 2018; Trueba et al., 2019). Recently, plant scientists have begun to pay renewed attention to the drought adaptations found in succulent plants (Heyduk et al., 2016; Males, 2017; Fradera-Soler et al., 2021; Leverett et al., 2021). Succulence is defined by the presence of water stores, in the leaf, stem and/or roots, which can be mobilized when a plant is dehydrated (Ogburn and Edwards, 2010). Typically, succulent tissues (i.e. the tissues responsible for water storage) arise due to the development of enlarged cells, either in the photosynthetic tissue (chlorenchyma), in a specialized achlorophyllous water-storage tissue (hydrenchyma), or a combination of the two (Eggli and Nyffeler, 2009; Borland et al., 2018; Heyduk, 2021; Leverett et al., 2022). If water stored in large cells can be mobilized during drought, succulent plants can dehydrate whilst maintaining water potentials (Ψ) at safe, stable levels. By buffering plant Ψ, succulence prevents a number of detrimental processes from occurring, such as the closing of stomata, the buckling of cells and the formation of emboli in the xylem (Brodribb et al., 2016; Vollenweider et al., 2016; Zhang et al., 2016; Henry et al., 2019). The benefits conferred by succulence have resulted in the succulent syndrome being found in plants across the globe, following adaptive radiations into the world’s arid and semi-arid ecosystems (Arakaki et al., 2011).

The adaptive benefits of succulence have recently drawn the attention of synthetic biologists, who have begun to recognize the potential this adaptation could have for food security and bioenergy in a drying world (Borland et al., 2009; Grace, 2019). Both modelling and field trials have assessed the value of growing succulent *Agave* and *Opuntia* in dry marginal and underused lands (Owen and Griffiths, 2014; Davis et al., 2017; Hartzell et al., 2021; Neupane et al., 2021). Furthermore, progress has been made to synthetically produce succulence in non-succulent species. The introduction of an exogenous transcription factor gene into *Arabidopsis thaliana* led to increased tissue succulence and higher water-use efficiency (Lim et al., 2018, 2020). These findings strongly suggest that bioengineering succulence has the potential to enhance drought resistance in crops. Whilst some work has been done to understand the genetic programs controlling the development of succulence (Heyduk, 2021), a great deal more research is needed if we are to fully utilize this adaptation in agricultural settings. In addition, we must appreciate every important trait that makes up the succulent syndrome. Beyond the genetic control of cell size, succulent species often exhibit a number of other co-adaptive traits, such as 3D vascular patterning, crassulacean acid metabolism (CAM) and waxy cuticles (Griffiths and Males, 2017). Cell walls have recently been postulated as an often-overlooked key component of the succulent syndrome (Ahl et al., 2019; Fradera-Soler et al., 2022), yet the precise mechanistic relevance of cell walls in succulent tissues remains largely speculative. In the present study, we analyse diverse succulent species and propose that cell wall biomechanics and biochemistry should be considered among the characteristic components of the succulent syndrome.

### Cell wall biomechanics in succulents

All plant cells are encased in a lattice-like structure, the cell wall (Popper et al., 2011). Primary, extensible cell walls are complex and dynamic systems composed largely of polysaccharides, polyphenols and certain types of glycoproteins (Carpita et al., 2015). When plant cells are hydrated, an osmotic gradient exists across the plasma membrane which results in water moving into the protoplasm (Beadle et al., 1993). This intake of water causes the plasma membrane to push against the cell wall, generating a positive pressure called turgor (*P*). The bulk modulus of cell wall elasticity (ε) relates to *P* according to the equation:

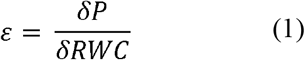

where relative water content (RWC) is the percentage of total water present in a tissue. Higher values of ε indicate greater cell wall rigidity and thus more resistance for the plasma membrane to push against, with changes in RWC resulting in large changes in *P*. Conversely, when ε is low and cell walls are highly elastic, changes to RWC have a lower impact on *P*, because cell walls can stretch and provide less resistance.

For succulent plants, ε has the potential to affect the point at which turgor is lost. As plant tissues dehydrate, Ψ falls, which results in a linear drop in *P* (Beadle et al., 1993). Eventually, Ψ falls to a point where *P* = 0, meaning there has been a total loss of turgor. When this turgor loss point (TLP_Ψ_) has been reached, leaves will typically wilt and cells will begin to experience damage (Trueba et al., 2019). Consequently, it is beneficial for plants to avoid reaching their TLP_Ψ_ (Kunert et al., 2021). Bartlett et al. (2012) found that the TLP_Ψ_ can be estimated by:

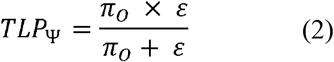

where π_O_ is the osmotic potential of fully hydrated tissues (a more negative π_O_ corresponds to a higher concentration of osmotically active solutes). Modifying ε or π_O_ are named elastic and osmotic adjustment, respectively, and can be used to alter the TLP_Ψ_ in order to allow cells to maintain turgor at more negative water potentials. Lower ε could result in cell walls capable of changing shape and folding as the protoplasm within shrinks (Ahl et al., 2019; Fradera-Soler et al., 2022). This would prevent the catastrophic disruption of the membrane-wall continuum and other forms of irreversible damage due to mechanical stress which occur when the TLP_Ψ_ is reached. However, studies of non-succulent species have found that ε is generally so high that changes to this trait are inconsequential for the TLP_Ψ_ (Bartlett et al., 2012). Put differently, in non-succulent species, cell walls are quite rigid, which means that even substantial changes to their elastic properties will not affect their TLP_Ψ_. This can be visualized by considering **Fig. 1A**. If π_O_ is held constant and ε is allowed to vary, the TLP_Ψ_ can be simulated using **Eq. 2**. This simulation forms a curve, and in non-succulent tissues the true value of ε intersects at the flat portion of the curve. Consequently, the phenotypic space inhabited by non-succulent species is one where changes to ε have no effect on the TLP_Ψ_.

**Fig. 1:**
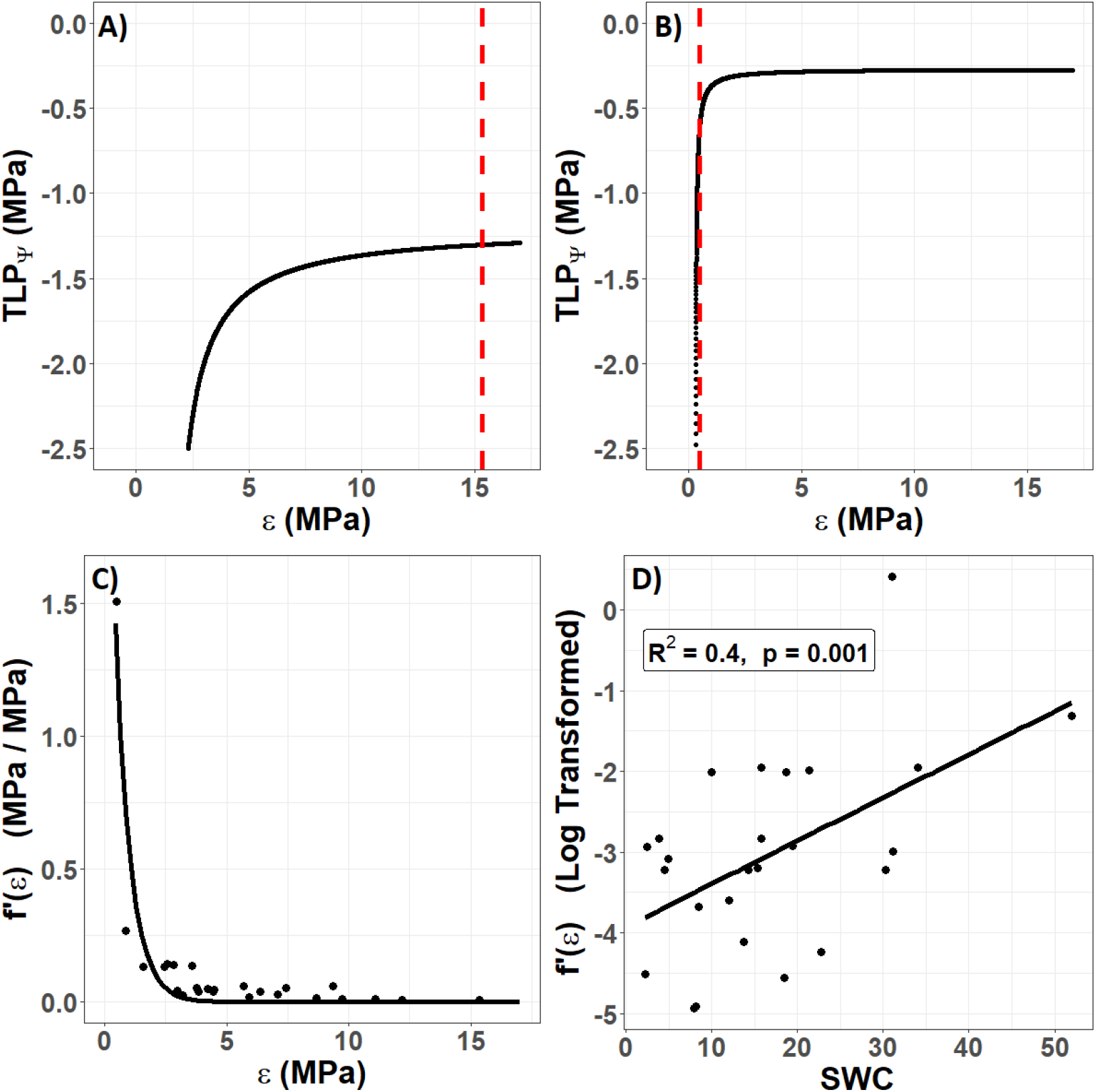
The unique role of cell wall biomechanics in succulent species. Using data published by Ogburn and Edwards (2012), the TLP_Ψ_ was simulated according to **Eq. 2** by holding π_O_ constant for each species and varying ε. (A) In non-succulent species, such as *Calandrinia colchaguensis*, the true value of ε (dashed line) intersects at the flat portion of the curve. Hence, changes to ε have little to no effect on the TLP_Ψ_. (B) In some succulent species, such as *Grahamia bracteata*, the true value of ε intersects at the curved potion of the line, meaning changes to ε effects the TLP_Ψ_. A quantitative estimate of the extent to which changing ε effects the TLP_Ψ_ was generated by finding the derivative of the curve at the point where the dashed line intersects [f’(ε)]. (C) In the Caryophyllales, lower values of ε result in exponentially higher values of f’(ε). An exponential curve still fit these data well when the species with the highest f’(e) value was removed (data not shown). (D) Saturated water content (SWC) correlates with the f’(ε), after this value has been log transformed.

The primary cell walls in succulent tissues are generally very thin and elastic (Goldstein et al., 1991; Ogburn and Edwards, 2010). Thus, the true value of ε for succulent species more often falls on the curved portion of the line (**Fig. 1B**). This means that for many succulent tissues, changes to cell wall biomechanics through elastic adjustment would have a much more substantial effect on the TLP_Ψ_ than in non-succulent plants. We sought to quantify this effect of ε on TLP_Ψ_ by repeating the simulation in **Figs. 1A-B** for several species. Ogburn and Edwards (2012) studied the relationship between parameters derived from pressure–volume curves and measures of succulence in the Caryophyllales, an angiosperm order comprising many succulent-rich groups with a broad range of tissue succulence. Using their published data, π_O_ was held constant for each species and ε was allowed to vary in order to simulate the TLP_Ψ_ according to **Eq. 2**. Then, for each species, we found the derivative of the curve, at the true value of ε (i.e. where the dashed line intersects the curve). This derivative, f’(ε), is a quantitative estimate of the extent to which changing ε affects the TLP_Ψ_. As ε values become very low in highly succulent species, f’(ε) becomes exponentially higher (**Fig. 1C**). Finally, we explored the relationship between f’(ε) and saturated water content (SWC), as the latter has been shown to be a powerful metric to quantify succulence in the Caryophyllales (Ogburn and Edwards, 2012). Log-transformed estimates of f’(ε) correlated significantly with SWC, using a linear regression model (**Fig. 1D**).

Together, our data show that unlike non-succulent species, succulent plants occupy a phenotypic space in which increases in cell wall elasticity during drought (i.e. elastic adjustment) can result in substantial decreases in TLP_Ψ_. Furthermore, once a succulent species moves into this phenotypic space, decreasing ε has an exponential effect on f’(ε), so that alterations to cell wall biomechanics become an increasingly efficient means of controlling the TLP_Ψ_. This agrees with the recently observed drought-induced modifications of pectic polysaccharides in hydrenchyma cell walls of *Aloe* (Ahl et al., 2019), which are believed to be a form of elastic adjustment that allows them to fold as cells shrink during dehydration (**Fig. 3**).

### Cell wall biochemistry in succulents

One way to assess the biochemical composition of cell walls is to investigate the extracellular glycome, which encompasses the entirety of extracellular carbohydrates in a tissue, organ or plant, and the majority of which corresponds to the cell wall. Characterizing glycomic profiles across different plant species can indicate which cell wall components have been favoured under different environmental conditions. Whilst the glycomes of some economically important succulent taxa, such as *Agave, Aloe* and *Opuntia*, have recently been analysed (Ginestra et al., 2009; Li et al., 2014; Ahl et al., 2018; Jones et al., 2020), little has been done to compare the cell wall composition of other distantly related succulent species. Hence, we sought to test the hypothesis that the extracellular glycome of phylogenetically diverse succulent species will exhibit some differences from those of non-succulents, so that a common ‘succulent glycome’ emerges. To this end, we sampled leaf material from 10 species with succulent leaves and 10 with non-succulent leaves, representing diverse lineages within the angiosperms (**Table 1**). Using the succulence index (SI) from Ogburn and Edwards (2010) as a proxy for the degree of succulence (see **Supplementary material**), these two groups differed significantly (*P* < 0.01) (**Fig. 2**). We used comprehensive microarray polymer profiling (CoMPP) to estimate and compare the relative polysaccharide contents of leaves from these species (see **Supplementary material**) (Moller et al., 2007; Ahl et al., 2018). We used whole leaves for comparability across species, assuming that mesophyll tissues would dominate the results. In the current study we used three extraction steps: water (targeting soluble unbound or loosely bound polysaccharides), CDTA (targeting primarily pectins) and NaOH (targeting primarily hemicelluloses). CoMPP relies on antibody-based molecular probes, so we used 49 monoclonal antibodies (mAbs) to target the majority of known cell wall polymer motifs (Moller et al., 2007; Rydahl et al., 2018) (**Tables S1–S3**). No representatives of commelinid monocots were included, given that their type-II cell wall biochemistry is particularly distinct from that of the rest of angiosperms (Carpita et al., 2015).

**Table 1.**
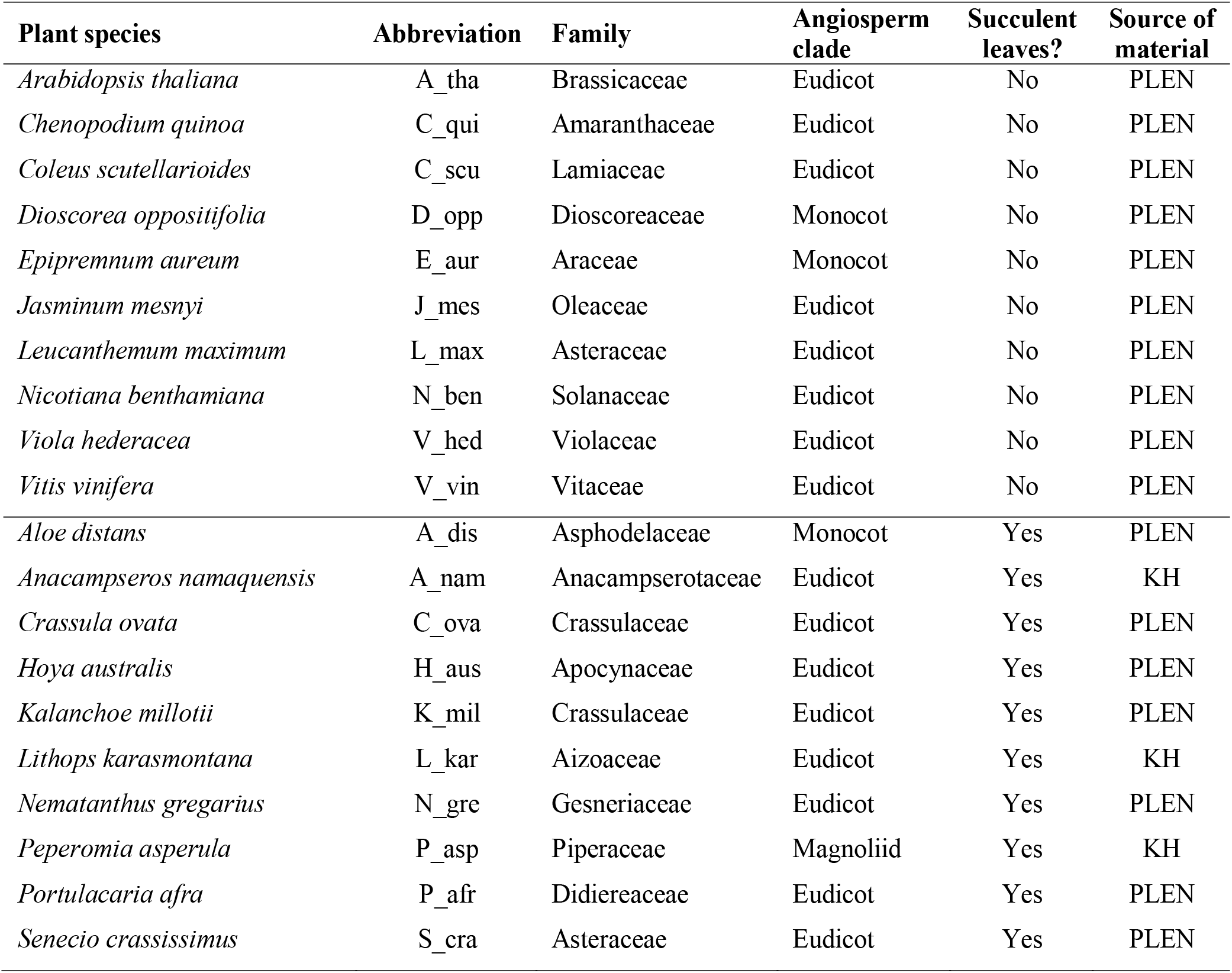
Angiosperm species used in this study. Abbreviations for the source of plant material: Department of Plant and Environmental Sciences, University of Copenhagen (PLEN), Kakteen-Haage nursery (KH).

| Plant species | Abbreviation | Family | Angiosperm clade | Succulent leaves? | Source of material |
| --- | --- | --- | --- | --- | --- |
| *Arabidopsis thaliana* | A_tha | Brassicaceae | Eudicot | No | PLEN |
| *Chenopodium quinoa* | C_qui | Amaranthaceae | Eudicot | No | PLEN |
| *Coleus scutellarioides* | C_scu | Lamiaceae | Eudicot | No | PLEN |
| *Dioscorea oppositifolia* | D_opp | Dioscoreaceae | Monocot | No | PLEN |
| *Epipremnum aureum* | E_aur | Araceae | Monocot | No | PLEN |
| *Jasminum mesnyi* | J_mes | Oleaceae | Eudicot | No | PLEN |
| *Leucanthemum maximum* | L_max | Asteraceae | Eudicot | No | PLEN |
| *Nicotiana benthamiana* | N_ben | Solanaceae | Eudicot | No | PLEN |
| *Viola hederacea* | V_hed | Violaceae | Eudicot | No | PLEN |
| *Vitis vinifera* | V_vin | Vitaceae | Eudicot | No | PLEN |
| *Aloe distans* | A_dis | Asphodelaceae | Monocot | Yes | PLEN |
| *Anacampseros namaquensis* | A_nam | Anacampserotaceae | Eudicot | Yes | KH |
| *Crassula ovata* | C_ova | Crassulaceae | Eudicot | Yes | PLEN |
| *Hoya australis* | H_aus | Apocynaceae | Eudicot | Yes | PLEN |
| *Kalanchoe millotii* | K_mil | Crassulaceae | Eudicot | Yes | PLEN |
| *Lithops karasmontana* | L_kar | Aizoaceae | Eudicot | Yes | KH |
| *Nematanthus gregarius* | N_gre | Gesneriaceae | Eudicot | Yes | PLEN |
| *Peperomia asperula* | P_asp | Piperaceae | Magnoliid | Yes | KH |
| *Portulacaria afra* | P_afr | Didiereaceae | Eudicot | Yes | PLEN |
| *Senecio crassissimus* | S_cra | Asteraceae | Eudicot | Yes | PLEN |

**Fig. 2:**
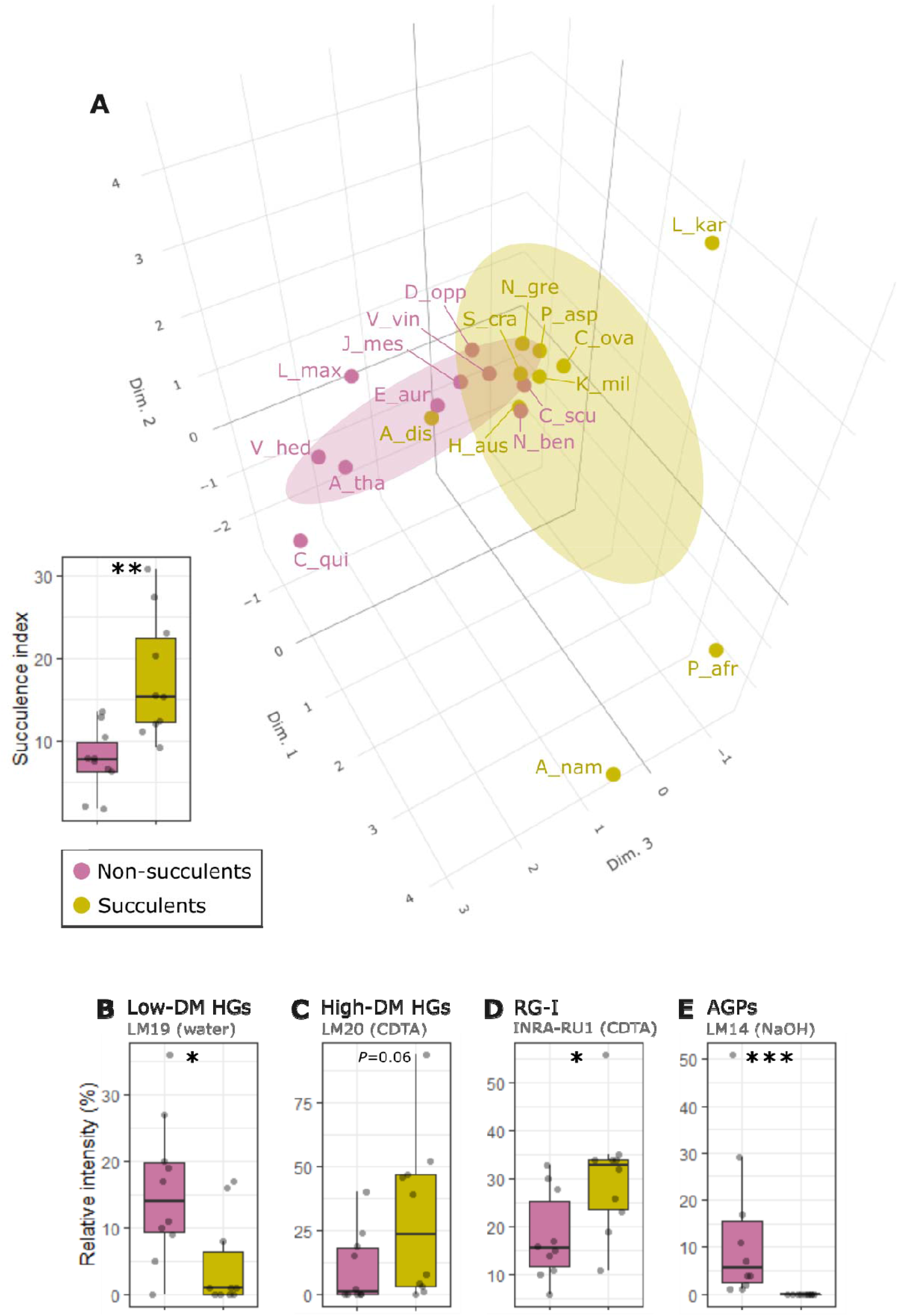
The succulent glycome: extracellular glycomic differences between succulents and non-succulents. (A) 3D score plot of the first three MFA dimensions (17%, 12.1% and 10.8% of total variance respectively) of glycomic data from 10 leaf succulents and 10 non-succulents (see **Table 1** for abbreviations), with concentration ellipsoids for each group. Succulents and non-succulents occupy distinct phenotypic spaces, particularly along dimension 3. On the left, boxplot of succulence index (SI) values for all the species; the two groups differ significantly. (B–E) Selection of antibodies showing significant differences between succulents and non-succulents for (B–D) pectins and (E) glycoproteins. Significant differences between the two groups, assessed using either Welch’s *t*-test (if both are normally distributed) or Wilcoxon test, are indicated by asterisks.

CoMPP results were analysed using multiple factor analysis (MFA) (see **Supplementary material**), which indicated that succulent species occupy a distinct phenotypic space different from non-succulent species (**Fig. 2A**). Of particular note is MFA dimension 3, along which succulents and non-succulents differed significantly (*P* < 0.01) and was driven mostly by glycoprotein- and pectin-targeting mAbs (**Fig. S1**). Three succulent species were “pulling” along dimension 1 and fell far from the main cluster, but even when omitting these three outliers from the MFA, the results still showed a significant difference between succulents and non-succulents (**Fig. S2**). We observed a higher signal for homogalacturonans (HGs) with a high degree of methyl-esterification (DM) in succulents (**Fig. S4**), which influences the nature of pectin gels (Willats et al., 2001; Hocq et al., 2017; Wormit and Usadel, 2018) and increases cell wall elasticity (Peaucelle et al., 2011; Levesque-Tremblay et al., 2015; Bidhendi and Geitmann, 2016). In contrast, non-succulents had a higher signal for low-DM HGs, which may indicate stiffer cell walls. Furthermore, we observed a higher signal for rhamnogalacturonan I (RG-I) backbones in succulents compared to non-succulents (**Fig. S5**). RG-I and its side chains (i.e. arabinans, galactans and/or arabinogalactans) have been postulated as cell wall plasticizers, which is a crucial feature for cells undergoing structural wall changes during dehydration and rehydration (Harholt et al., 2010; Moore et al., 2013; Kaczmarska et al., 2022). Together, MFA of the CoMPP results suggests that fundamental differences exist between the cell wall composition of diverse succulent and non-succulent species (**Fig. 2B**). The three outlying succulent species (*Anacampseros namaquensis, Lithops karasmontana* and *Portulacaria afra*) belong to the core Caryophyllales, and two of them (*A. namaquensis* and *P. afra*) to suborder Portulacineae. These taxa showed remarkably high signal for RG-I as well as soluble glucuronoxylans (**Figs. S5, S6**), which most likely reflects the presence of highly hydrophilic apoplastic mucilage in succulents in the Caryophyllales, particularly those in the Portulacineae (**Fig. 3**) (Hernandes-Lopes et al., 2016; Cole, 2020).

**Fig. 3:**
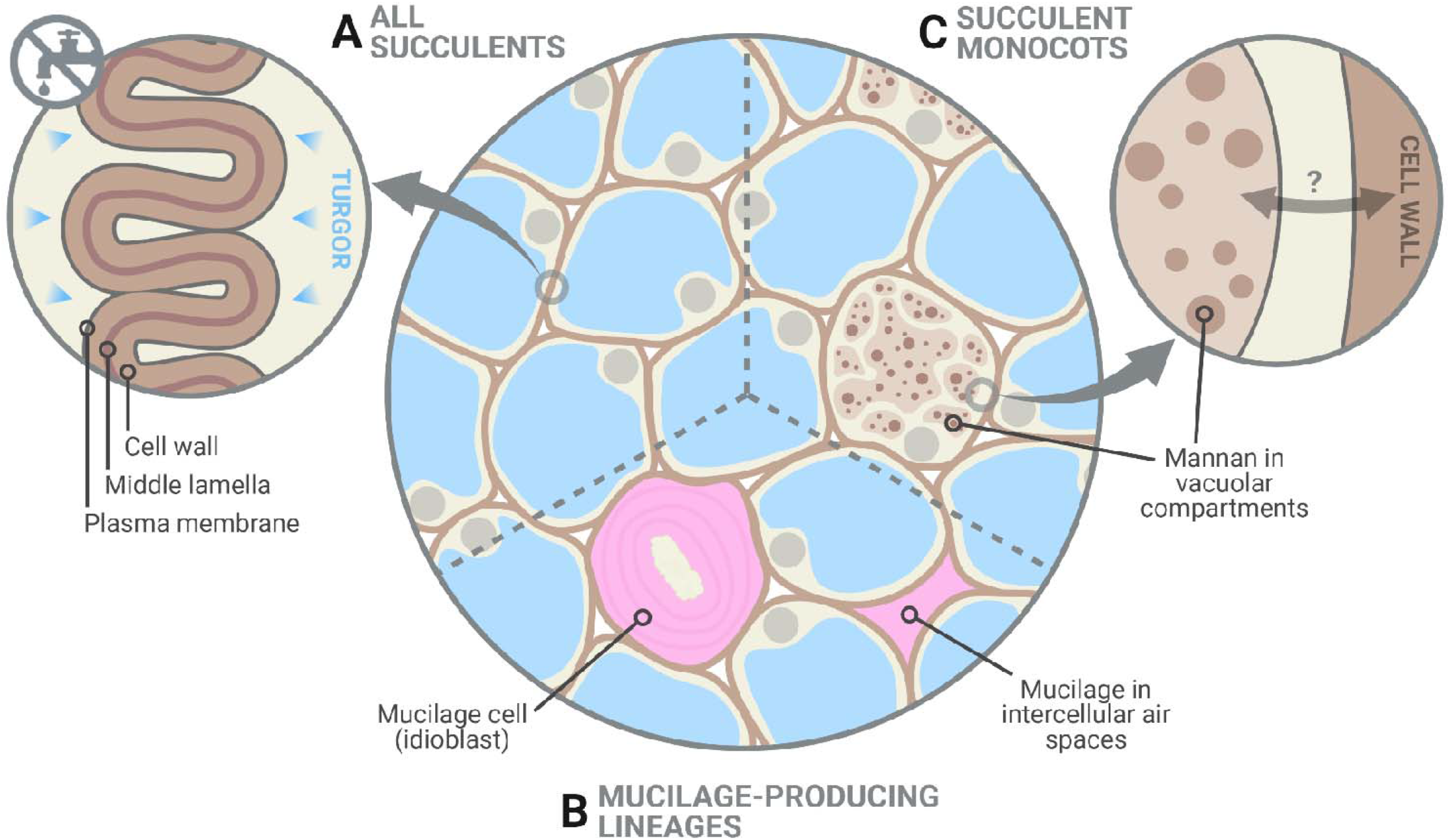
Glycomic diversity among succulent plants. (A) Succulent tissues have thin and highly elastic cell walls and, as shown in this study, elastic adjustment through cell wall remodelling likely plays a crucial role in preventing turgor loss during dehydration. Despite the clear differences between succulents and non-succulents, we also noted considerable glycomic diversity among succulents. (B) Mucilage-producing succulent lineages, mostly those in the Caryophyllales and particularly in the Portulacineae, accumulate pectin-rich mucilage in the periplasmic space of mucilage cells and/or in intercellular spaces, which boosts their water-storage capacity (Mauseth, 2005; Ogburn and Edwards, 2009). (C) Storage mannans can be found in vegetative tissues of many monocot lineages, often stored within vacuolar compartments. In succulent monocots, mobilization of these mannans may be part of the drought response, as seen in *Aloe* (Ahl et al., 2019), in which cell wall-associated mannan is remobilized into the protoplasm. However, the dynamics between cell-wall associated and vacuolar mannans in monocots remain largely unexplored. Created with BioRender.com.

In addition to MFA, we used a random forest (RF) algorithm to determine whether extracellular glycomic profiles can be used to predict if a species is succulent or non-succulent (see **Supplementary material**). Based on CoMPP data alone, the RF algorithm was able to classify species in their respective categories with 90% accuracy. The variable importance plot from the RF algorithm identified several cell wall components driving this classification (**Fig. S3**), namely arabinogalactan proteins (AGPs), xylans, low-DM HGs and RG-I (incl. arabinan and galactan side chains). For instance, we observed drastically lower levels of xylans in the succulents studied, compared to the non-succulent species (**Fig. S6**). However, xylans are often found in lignified support tissues (Zhong et al., 2013), and small-stature succulent species such as the ones we studied generally lack these tissues, relying primarily on turgor for support (Niklas, 1992; Gibson, 1996; Bobich and North, 2009). Thus, such differences may not hold for larger succulents. An interesting observation concerns AGPs, a notoriously complex group of cell wall glycoproteins with many suggested functions, yet their precise mode of action is still uncertain (Seifert and Roberts, 2007; Ellis et al., 2010; Silva et al., 2020). Two mAbs targeting AGPs (LM14 and MAC207) did not yield any signal among succulents, despite being present in most non-succulent species tested (**Fig. S8**). Previous studies suggest that these two antibodies recognize the same or structurally related epitopes (Jackson et al., 2012; Marzec et al., 2015; Yan et al., 2015), which often exhibit a broad distribution across tissues (Amsbury et al., 2016; Wu et al., 2017; Leszczuk et al., 2019). Our results suggest that these epitopes are absent in some plant lineages, such as the Lamiids (*Coleus, Jasminum, Nicotiana*), in agreement with previous studies (Moller et al., 2008).

However, our results show that succulent representatives lack LM14 and MAC207 signals, even within lineages that are known to possess these epitopes. For example, succulent species from the Caryophylalles (*Anacampseros, Lithops* and *Portulacaria*) lacked these epitopes, whereas the non-succulent *Chenopodium quinoa* did not. Likewise, in the Asteraceae these epitopes were absent in the succulent species *Senecio crassissimus*, but present in the non-succulent *Leucanthemum maximum*. In contrast, other AGP-targeting mAbs showed comparable or slightly higher levels in succulents, likely reflecting the diversity of AGPs and their numerous alleged functions. Periplasmic AGPs for instance have been postulated as stabilizers of the membrane-cell wall continuum, and may also act as cell wall plasticizers when they are released from their membrane anchors (Gens et al., 2000; Knox, 2006; Lamport et al., 2006; Liu et al., 2015). The striking differences in signal intensity of the AGP-targeting mAbs we used warrant further exploration into the specific epitopes that they recognize and their functions.

The mobilization of soluble mannans has been suggested as a general drought response among succulents, based on studies of succulent leaves of *Aloe* and succulent-like storage organs of orchids and monocot geophytes (Ranwala and Miller, 2008; Wang et al., 2008; Chua et al., 2013; Ahl et al., 2019). However, our CoMPP data showed no clear difference between the mannans of succulents and non-succulents (**Fig. S7**). Instead, the few species with remarkably high signal for loosely bound, soluble mannans correspond to the three non-commelinid monocot species included in this study: *Aloe distans* (leaf succulent), *Dioscorea oppositifolia* and *Epipremnum aureum* (non-succulents). Among angiosperms, the presence of storage mannans in vegetative tissues seems to be restricted to monocots, with mannans being stored in granular or highly hydrated mucilaginous form within vacuolar cell compartments (Meier and Reid, 1982; He et al., 2017). Soluble mannans may therefore be uniquely important to monocots, being repurposed for drought response in succulent monocots (e.g. Ahl et al., 2019; **Fig. 3**), and not a component of a more general succulent glycome.

## Conclusions and future directions

Cell wall biomechanics and biochemistry of succulent leaves exhibit distinct differences from non-succulent species. In non-succulent species, highly rigid cell walls prevent elastic adjustment from having a physiologically meaningful impact on the TLP_Ψ_ (Bartlett et al., 2012). However, many succulent species have highly elastic cell walls, and our modelling indicates that even slight increases in cell wall elasticity (i.e. decreases in ε) in these species can have a large exponential effect on the TLP_Ψ_. Therefore, succulent plants use elastic adjustment advantageously during dehydration to acclimate to declining Ψ. In addition to biomechanical differences, our glycomic data show several similarities across phylogenetically diverse succulent taxa, namely a higher degree of HG methyl-esterification and a greater abundance of RG-I. These biochemical differences likely contribute to the high elasticity in the cell walls of succulent organs, which in turn facilitates the folding process during dehydration (Fradera-Soler et al., 2022). Interestingly, some glycomic features seem to be restricted to certain succulent lineages, pointing to some glycomic diversity among succulent plants: succulent monocots may have co-opted soluble mannans for drought response, whereas succulents in the Caryophyllales contain pectin-rich apoplastic mucilage which boosts their water-storage capacity. Together, our data demonstrate that succulent plants occupy a unique phenotypic space regarding both cell wall biomechanics and biochemistry. We suggest that cell wall traits should be regarded as one of the core components of the adaptations that make up the succulent syndrome.

Looking forward, it will be valuable to explore cell wall biology among closely related succulent taxa and considering cell wall trait heterogeneity within succulent organs. Cell wall thickness and elasticity are known to differ between hydrenchyma and chlorenchyma in some succulent organs (Goldstein et al., 1991; Nobel, 2006; Leverett et al., 2022), but further examination of cell wall biomechanics and biochemistry is needed to fully understand how these traits aid in whole-plant survival during drought. Ultimately, further research is needed into the dynamic nature of cell walls in succulent plants and to determine whether cell wall traits are indeed regulated during drought. Besides high-throughput methods based on immune-profiling such as CoMPP, our understanding of cell wall composition, structure and assembly in succulents can also be advanced using visualization with fluorescent probes (Rydahl et al., 2018; Bidhendi et al., 2020), high-resolution microscopy techniques (Zhao et al., 2019; DeVree et al., 2021), and nuclear magnetic resonance (NMR) (Zhao et al., 2020).

## Supporting information

Raw data and Random Forest output

## Abbreviations and symbols

**Physiological parameters**

ε: Bulk modulus of cell wall elasticity
*P*: Turgor pressure
RWC: Relative water content
TLP_Ψ_: Turgor loss point, i.e. water potential at which turgor is lost
Ψ: Water potential
π_O_: Osmotic potential of tissue at full hydration

**Cell wall polymers**

AGP: Arabinogalactan protein
HG: Homogalacturonan
RG-I: Rhamnogalacturonan I
DM: Degree of methyl-esterification
DP: Degree of polymerization

## Acknowledgements

The authors would like to thank Matthew Ogburn and Erika Edwards for the data that was used in the biomechanical modelling (see Ogburn and Edwards, 2012). The authors would also like to thank Theodor E. Bolsterli, Morten L. Stephensen, Ouda Khammy, Davide Visintainer and Luu Trinh (PLEN, University of Copenhagen) and the Kakteen-Haage nursery (Erfurt, Germany) for providing plant material, and Sylwia Głazowska and Jeanett Hansen (PLEN, University of Copenhagen) for help and support during laboratory work.

This research was partially funded by Newcastle University’s R. B. Cook Scholarship.

This project has received funding from the European Union’s Horizon 2020 research and innovation programme under the Marie Skłodowska-Curie grant agreement No 801199.

