## Supplementary material for "Are cell wall traits a component of the succulent syndrome?": Methods

**Succulence index**

The succulence index (SI) is a dimensionless ratio between water content of a succulent organ and its dry weight. It was presented by Ogburn and Edwards (2010), derived from metrics used by von Willert et al. (1990, 1992). The SI was calculated as:

$$SI = \frac{Fresh weight at full hydration-Dry weight}{Dry weight}$$

For each taxon, 5 leaves were sampled a day after thorough watering, to ensure full hydration under near-natural conditions. The fresh weight at full hydration was obtained by weighing the leaves on an analytical balance (XS204, d=0.1 mg, Mettler Toledo) a day after thorough watering to ensure maximum hydration under near-natural conditions. Leaves were then dried in an oven at ~80°C for at least 3 days and reweighed to obtain the dry weight. SI values from the 5 replicates were averaged.

**Comprehensive microarray polymer profiling (CoMPP)**

***Polysaccharide extraction***

The CoMPP protocol followed in the current study is a modification of the one described by Moller et al. (2007), which in turn was optimized for succulent tissues by Ahl et al. (2018). The analysis was performed on alcohol-insoluble residue (AIR), which was obtained through the ‘70-CHCl_3_/MeOH L’ protocol (Fangel et al., 2021). 10 mg of AIR material were placed in tubes with a glass bead; two technical replicates were used for each taxon. A three-step sequential extraction series was used, with the following solvents: dH_2_O, 50 mM CDTA (trans-1,2-diaminocyclohexane-N,N,N′,N′-tetraacetic acid monohydrate; pH 7.5), and 4 M NaOH + 0.1% NaBH_4_. For each extraction step, a ratio of 600 µl of solvent to the initial 10 mg of AIR material was used, and the tubes were shaken at room temperature in a bead mill (TissueLyser II, Qiagen) at 27 s^−1^ for 2 min, and then at 6 s^−1^ for 2 h. Subsequent extractions were performed on the remaining pellet of the previous extraction step; the pellet was discarded after the last step. After each extraction, the tubes were centrifuged at 3,000 rpm for 10 min and the supernatants were carefully removed and transferred to new tubes, which were then stored at 4°C to minimize degradation. All extracts belonging to a specific extraction step is also known as fraction.

***Microarray printing***

The extracts were diluted in Arrayjet buffer adjusted to 15% v/v glycerol (83.6% dH_2_O, 15% glycerol, 1.4% Triton X (4.5 g/100 ml)) in order to avoid printing issues due to highly viscous samples among the succulent taxa. The extracts were then plated into 384-well microplates (Greiner Bio-One), and a four‐fold serial dilution was made for each extract in the microplates. The dilution series for all fractions was 1:4, 1:12, 1:36, 1:108. The microplates were printed on a 0.45‐μm nitrocellulose membrane (Whatman) using an Arrayjet Sprint (Arrayjet Ltd.) piezoelectric robotic printer; the printing was performed in duplicate on each microarray.

***Probing and development of microarrays***

The microarrays were first blocked under agitation at room temperature in PBS (pH 7.5) with 5% w/v skimmed milk powder (MPBS) for 1 h. A total of 49 monoclonal antibodies (mAbs), targeting different polysaccharides, were used to probe the microarrays as primary antibodies (Tables S2A–C); these mAbs were diluted in MPBS with different dilution ratios: either 1:10, 1:100 or 1:1000 (data not shown). The microarrays were incubated under agitation at room temperature in primary antibody solutions for 2 h. They were then washed in PBS three times and incubated under agitation at room temperature in secondary antibody solutions (goat-produced polyclonal antibodies conjugated to alkaline phosphatase, anti-rat or anti-mouse IgG) for 2 h. The microarrays were taken through three PBS washes and a final wash in dH_2_O. The microarrays were developed in the dark for 2–25 min in a solution composed of 7.5% BCIP (20 mg/ml; 5-bromo-4-chloro-3-indolyl-phosphate), 5.9% NBT (50 mg/ml; nitro blue tetrazolium chloride), and 86.6% alkaline phosphatase buffer (100 mM NaCl, 5 mM MgCl_2_, 100 mM Tris-HCl; pH 9.5). They were then moved to dH_2_O to stop the reaction and dried on filter paper.

***Quantification of microarrays***

The developed microarrays were scanned with a desktop scanner (Canon 8800) using ArchSoft PhotoStudio at a resolution of 2400 dpi. Quantification was performed on ImaGene 6.0 software (BioDiscovery) and Microsoft Office Excel (Microsoft). The intensity values of printed duplicates were averaged, and the values of the dilution series were subsequently averaged. For each of the three fractions separately, the highest intensity value was set to a value of 100, and the rest of the data was adjusted accordingly to represent a percentage value; no cut-off value was used in the current study. The results were presented in heatmap format.

**Statistical analyses**

Statistical analyses were performed on the CoMPP and SI data in RStudio (version 2022.02.0 Build 443; RStudio Team, 2022) using R (version 4.1.2; R Core Team, 2021).

***Multiple factor analysis***

Multiple factor analysis (MFA) was used as an integrative method to identify the main dimensions of variance in the whole CoMPP dataset (Pagès, 2014; Kassambara, 2017). MFA allows for variables to be structured in groups, and each group of variables is weighted during the analysis. Our variables were grouped according to the three CoMPP fractions (water, CDTA and NaOH), and variables with values of 0 across all species were omitted from the analysis. The analysis was performed using the R packages *factoextra* (Kassambara and Mundt, 2020) and *FactoMineR* (Lê *et al.*, 2008).

***Random forest analysis***

Random forest (RF) is a popular machine learning algorithm for data mining and pattern recognition in omics-scale data (Chen and Ishwaran, 2012; Touw *et al.*, 2013). A RF algorithm was used to classify the species into succulent or non-succulent based solely on the CoMPP dataset. The RF output also allowed us to identify the importance of the variables in this classification. The number of decision trees to be generated was set to 10^5^, and the number of variables used in the construction of each tree was set to 12. RF was performed using the R package *randomForest* (Liaw and Wiener, 2002).
