## Supplementary figures for "Are cell wall traits a component of the succulent syndrome?"

**Fig. S1:** Scatterplot matrix of the first three dimensions of MFA performed on 10 succulent (S) and 10 non-succulent species (NS). These two groups differ significantly in dimension 3 (Wilcoxon test, *P* < 0.01). On the right, contribution plots of the CoMPP fractions (water, CDTA and NaOH) to each of the MFA dimensions.

**Fig. S2:** Scatterplot matrix of the first three dimensions of MFA omitting the three succulent taxa belonging to the Portulacineae (*Anacampseros namaquensis*, *Lithops karasmontana* and *Portulacaria afra*). Succulents and non-succulents still differ significantly in dimension 1 (Wilcoxon test, *P* < 0.01).

**Fig. S3:** Variable importance plots from the Random Forest algorithm based on (A) the mean decrease of accuracy and (B) the mean decrease of the Gini impurity metric, showing the 30 most important variables (out of 98 CoMPP variables which yielded signal).

**Fig. S4:** Boxplots of succulents (S) versus non-succulents (NS) from a selection of HG-targeting antibodies. The y-axes represent relative intensity of signal within a specific CoMPP fraction. Significant differences between the two groups, assessed using either Welch's *t*-test (if both are normally distributed) or Wilcoxon test, are indicated by asterisks.

**Fig. S5:** Boxplots of succulents (S) versus non-succulents (NS) from a selection of RG-I-targeting antibodies. The y-axes represent relative intensity of signal within a specific CoMPP fraction. Significant differences between the two groups, assessed using either Welch's *t*-test (if both are normally distributed) or Wilcoxon test, are indicated by asterisks. Some outlying taxa have been labelled (see **Table S1** for abbreviations).

**Fig. S6:** Boxplots of succulents (S) versus non-succulents (NS) from a selection of xylan-targeting antibodies. The y-axes represent relative intensity of signal within a specific CoMPP fraction. Significant differences between the two groups, assessed using either Welch's *t*-test (if both are normally distributed) or Wilcoxon test, are indicated by asterisks. Some outlying taxa have been labelled (see **Table S1** for abbreviations).

**Fig. S7:** Boxplots of succulents (S) versus non-succulents (NS) from a selection of mannan-targeting antibodies. The y-axes represent relative intensity of signal within a specific CoMPP fraction. Some outlying taxa have been labelled (see **Table S1** for abbreviations).

**Fig. S8:** Boxplots of succulents (S) versus non-succulents (NS) from a selection of AGP-targeting antibodies. The y-axes represent relative intensity of signal within a specific CoMPP fraction. Significant differences between the two groups, assessed using either Welch's *t*-test (if both are normally distributed) or Wilcoxon test, are indicated by asterisks.

Figure S1

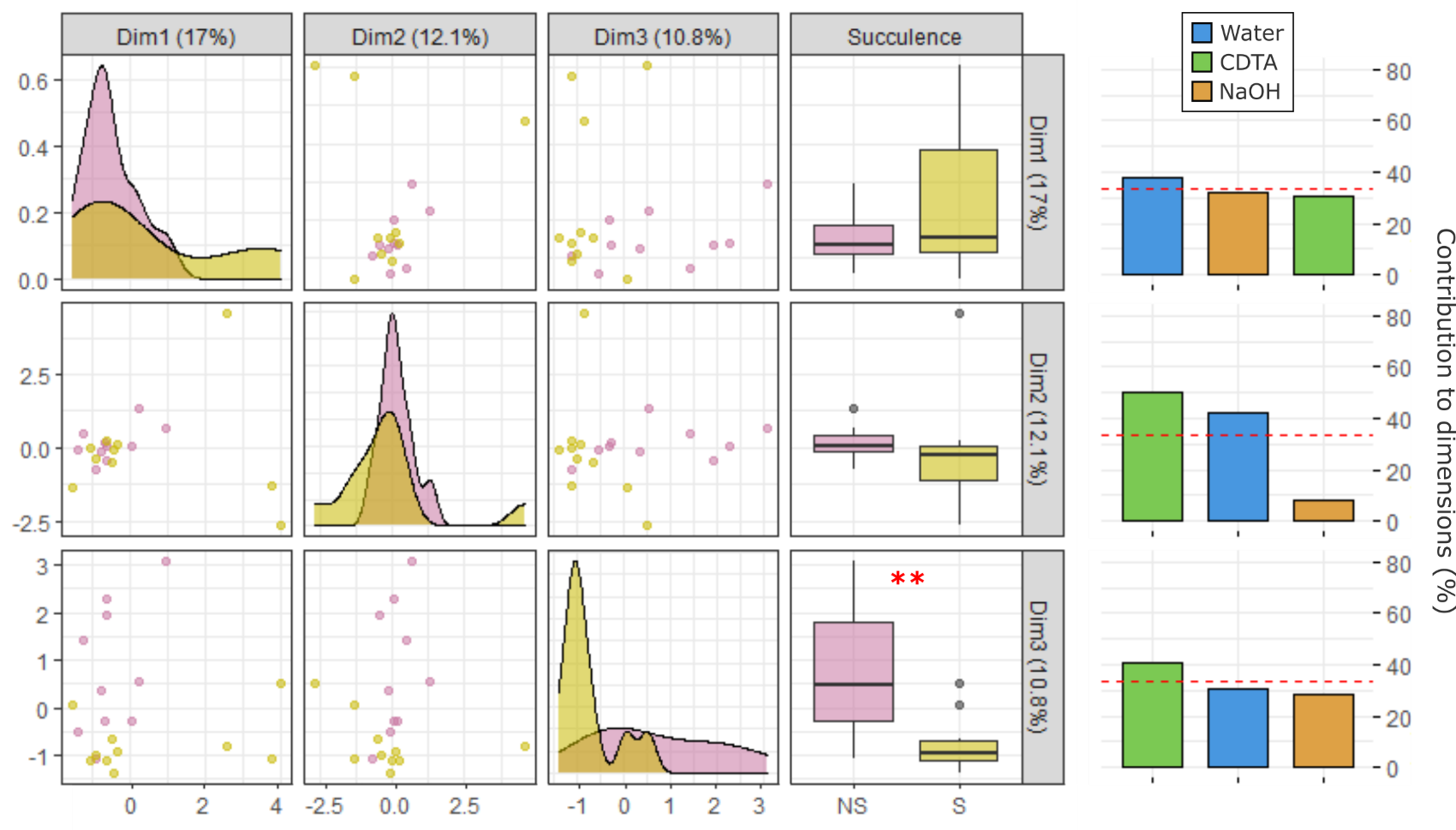

Figure S2

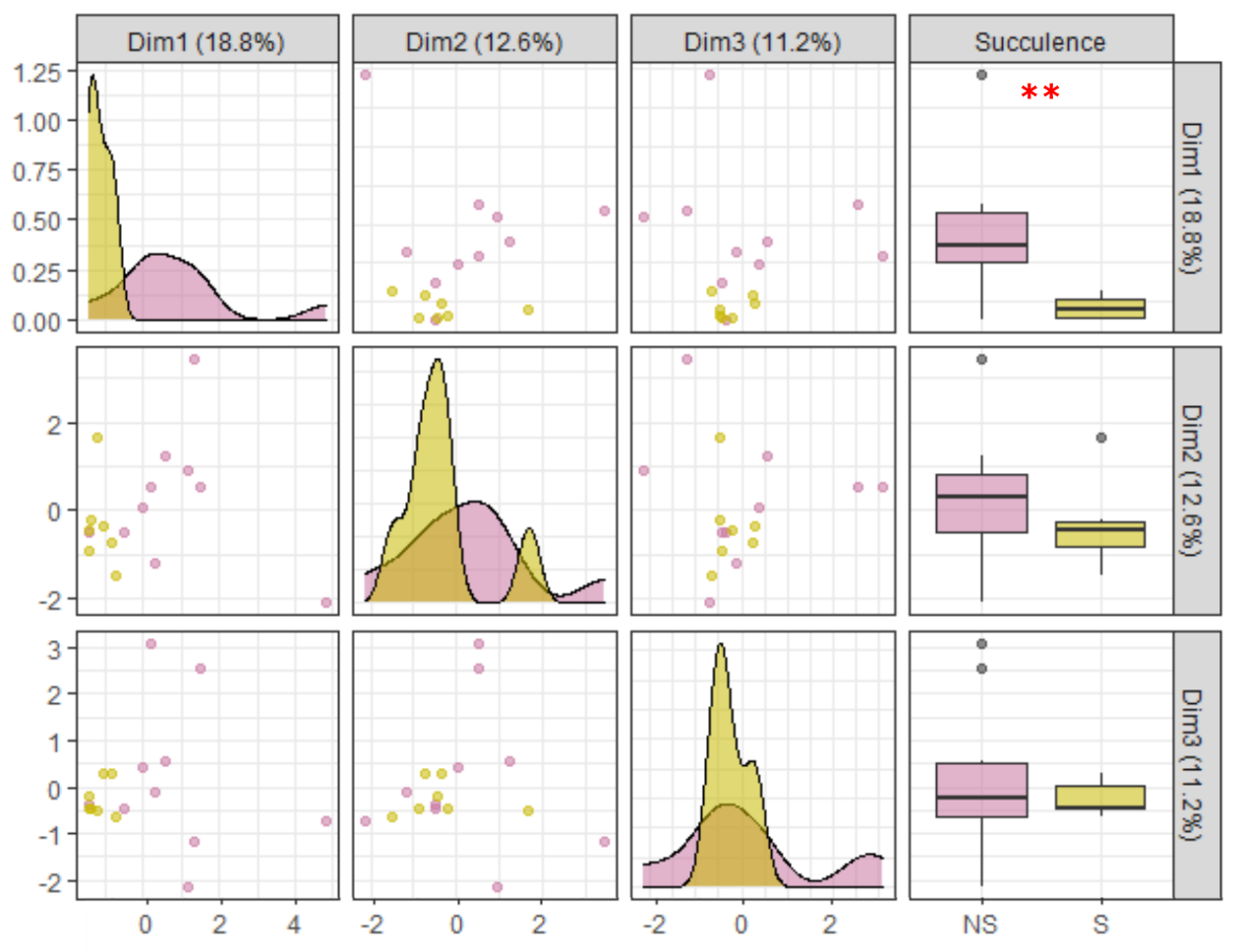

Figure S3

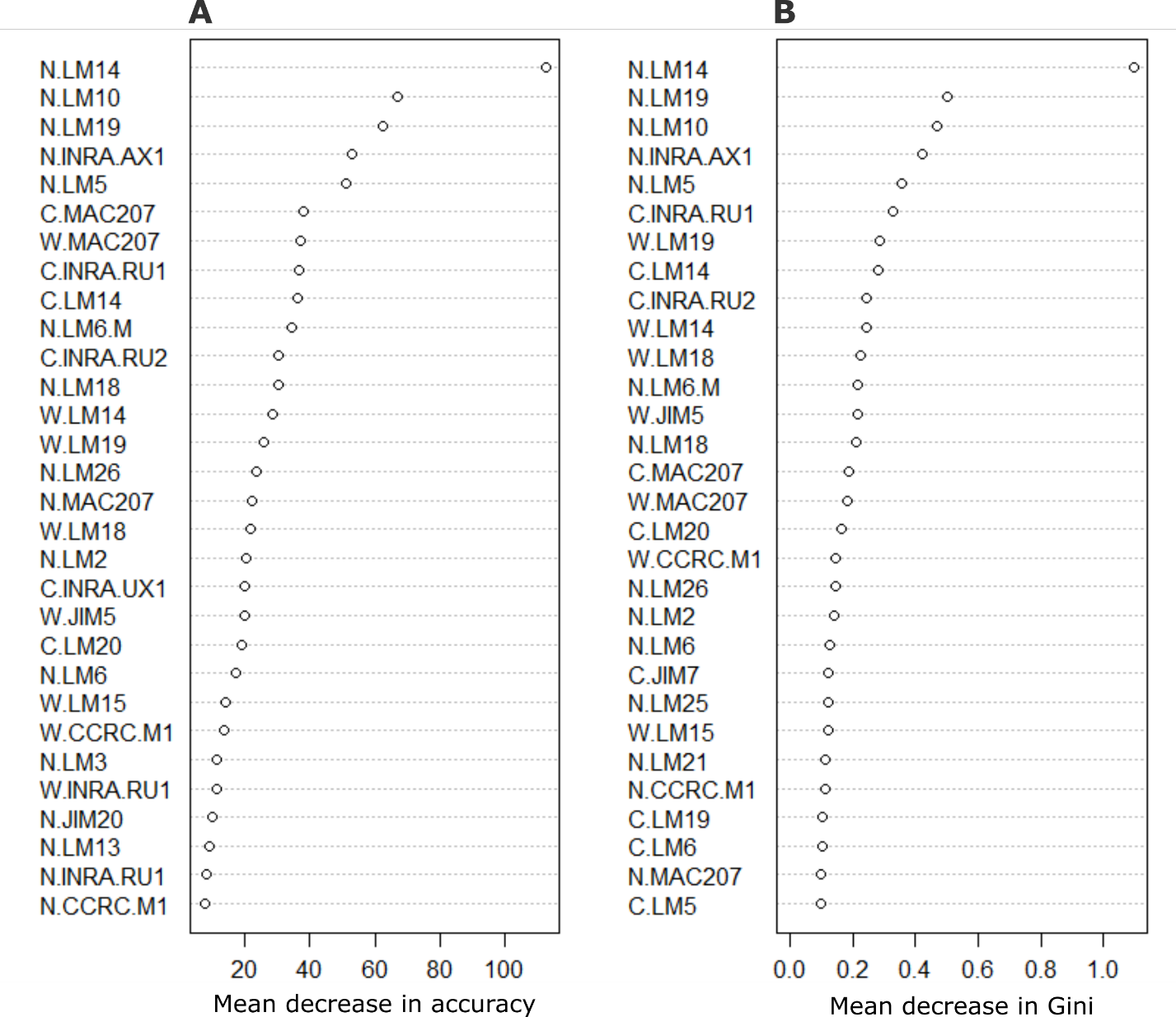

Figure S4

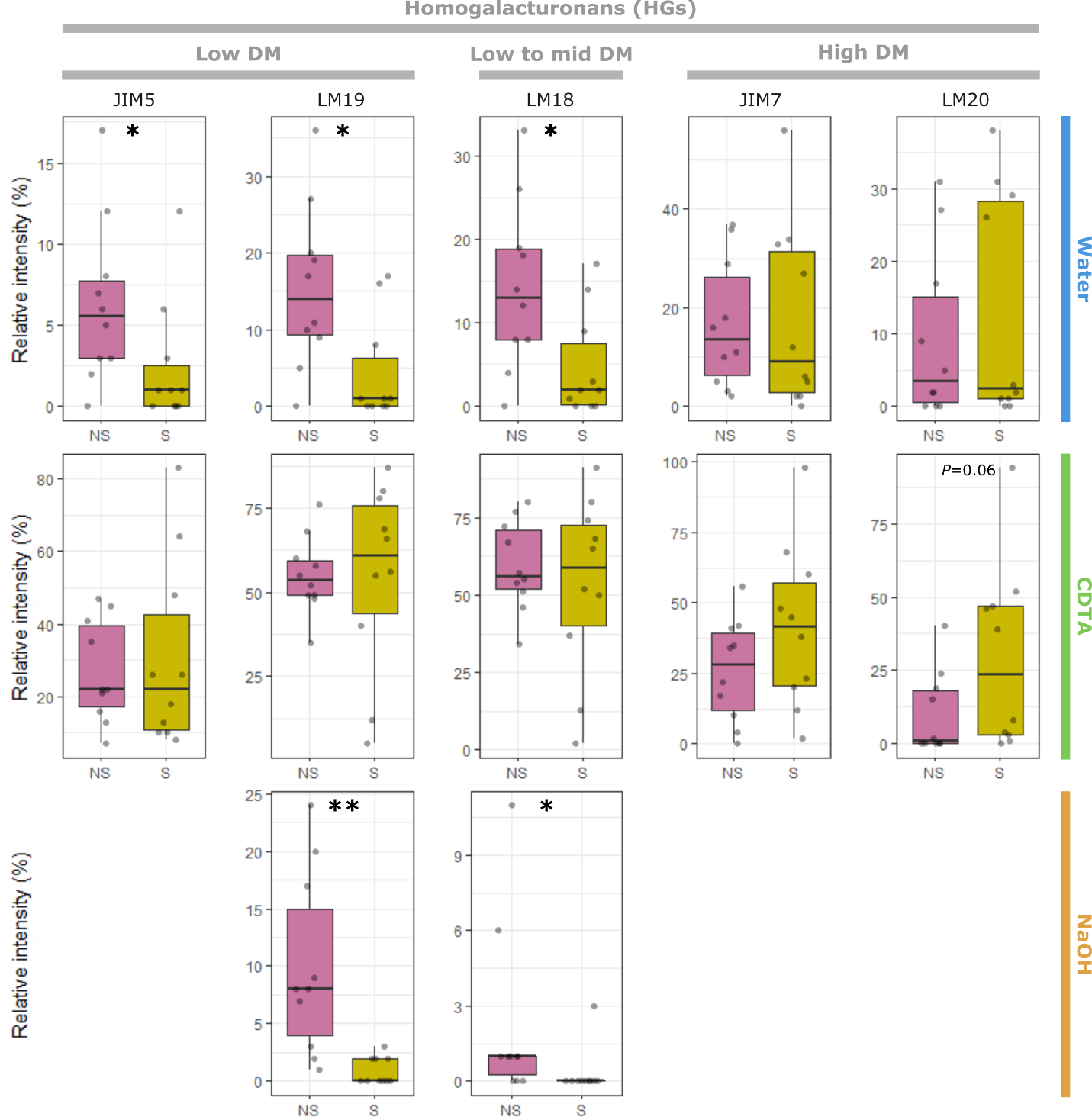

Figure S5

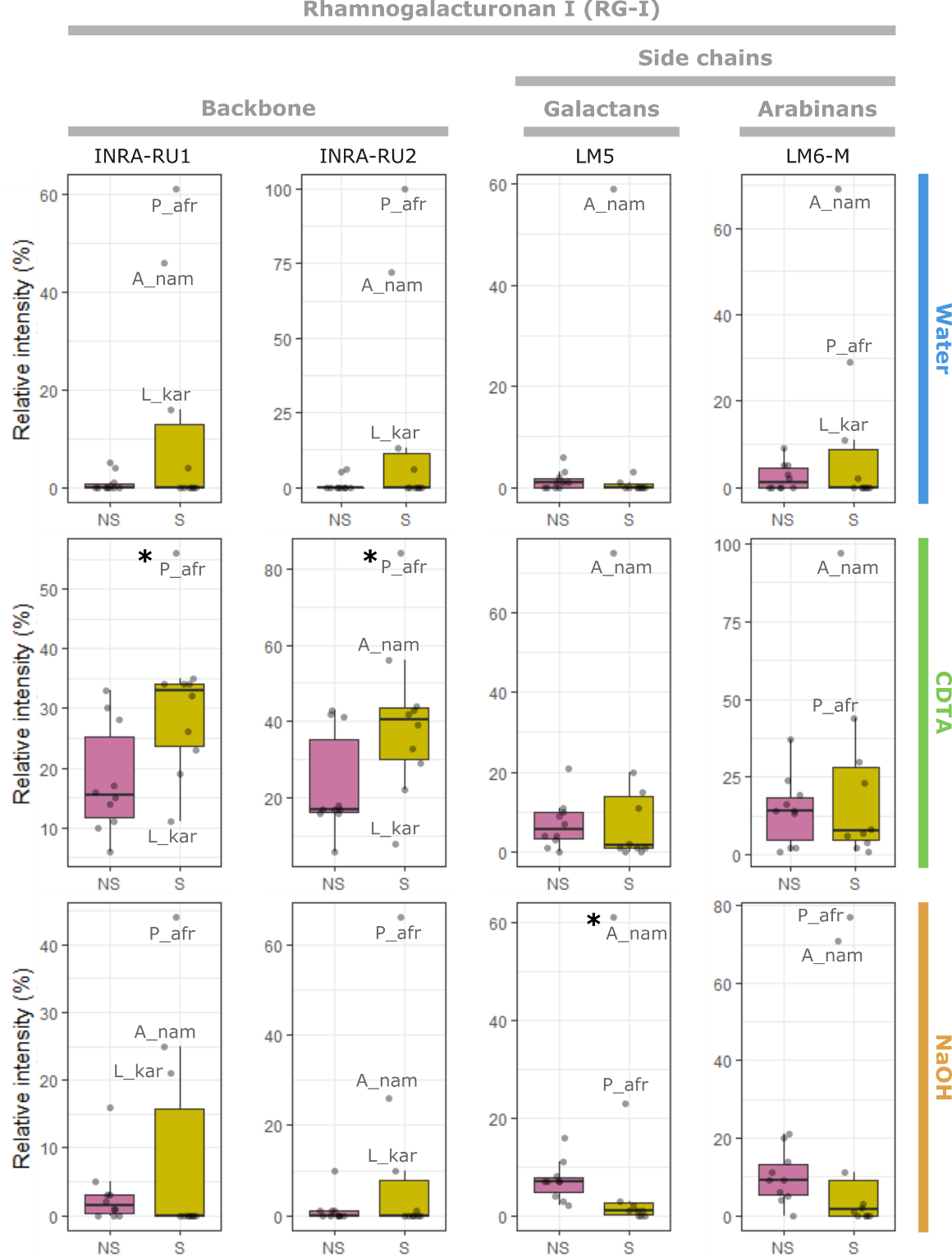

Figure S6

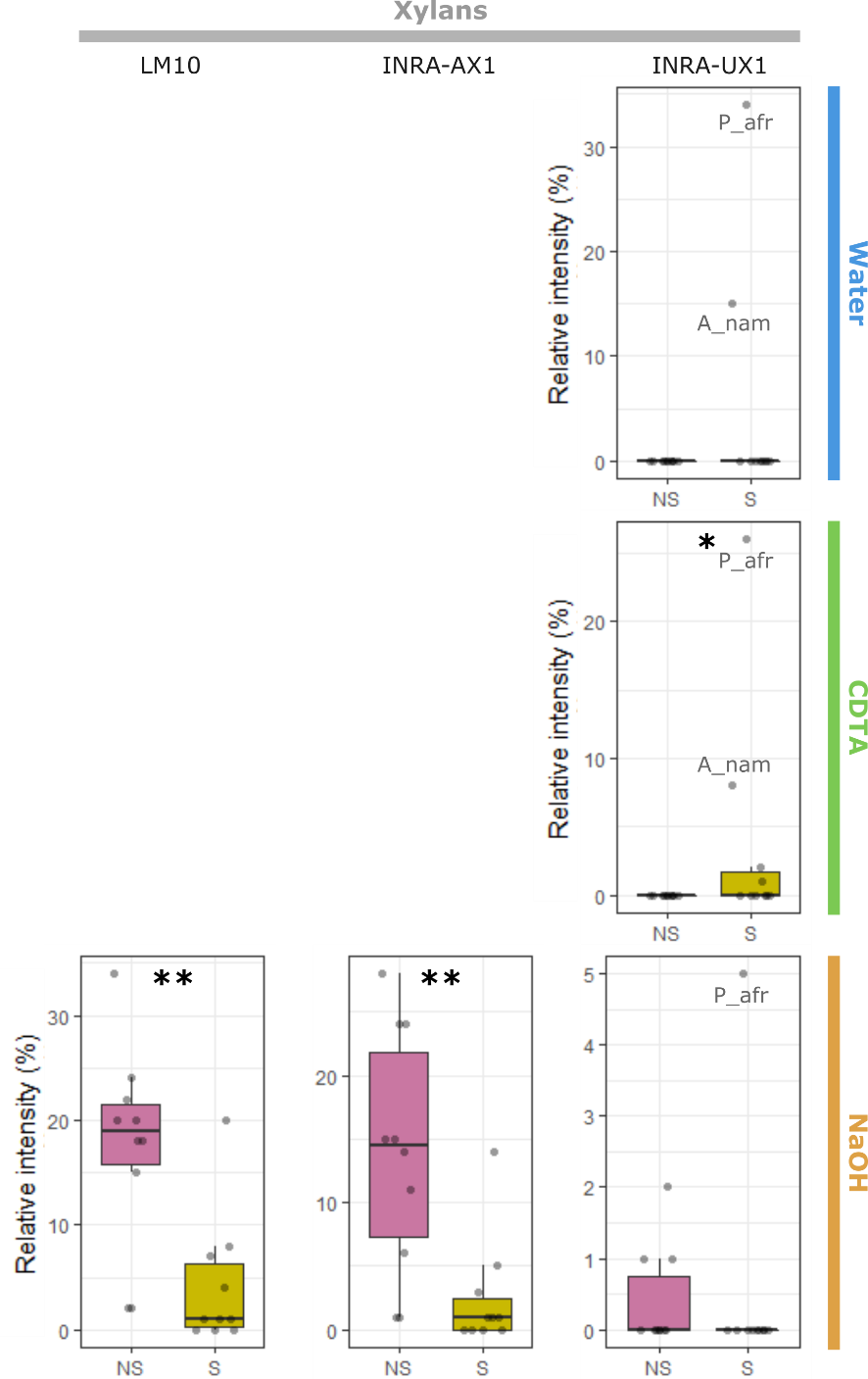

Figure S7

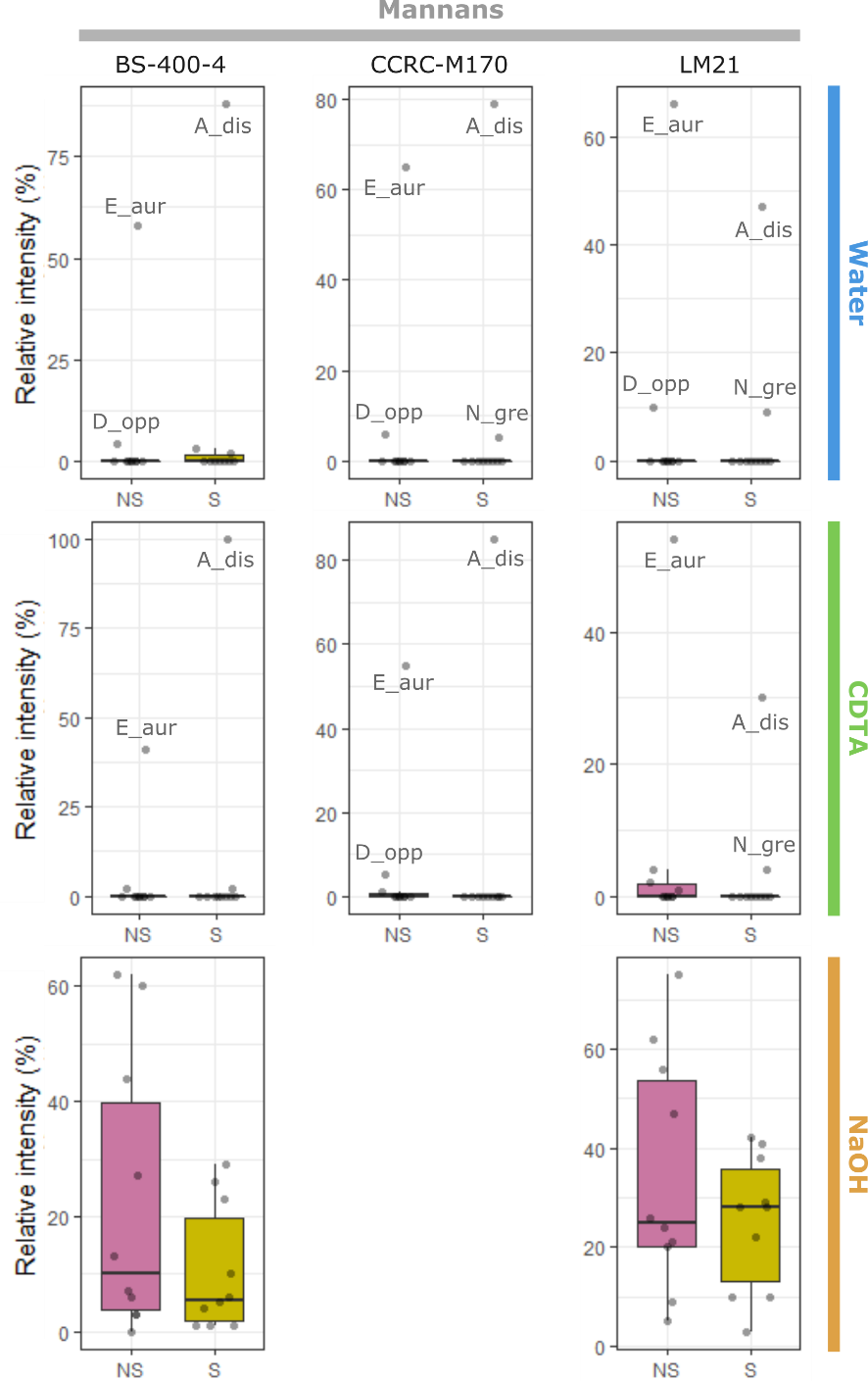

Figure S8

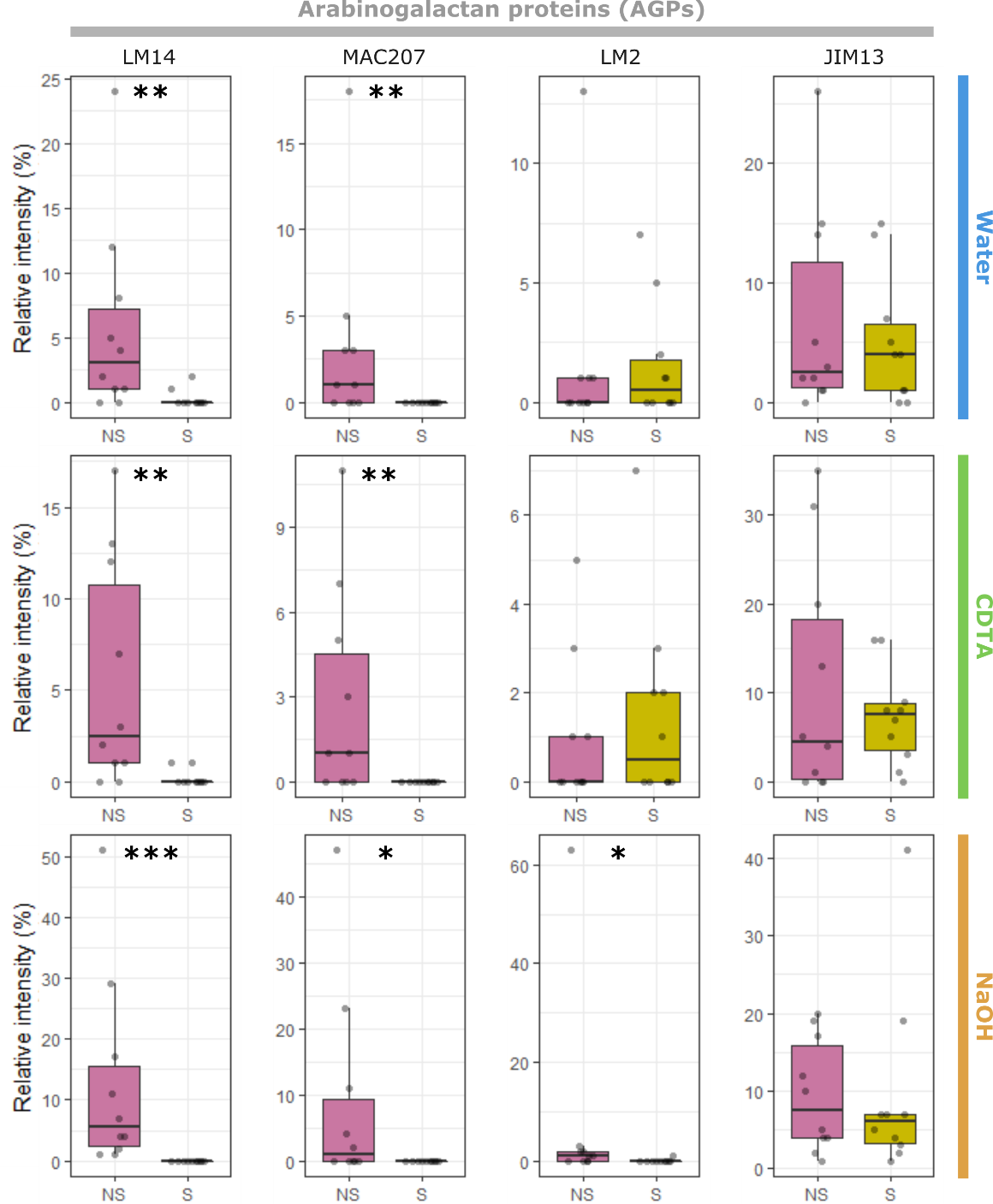
