## Supplementary tables for "Are cell wall traits a component of the succulent syndrome?"

**Table S1.** List of primary mAbs targeting pectins. Adapted from Rydahl *et al.* (2018) and references therein, unless otherwise indicated.

| **mAb** | **Antigen / Epitope** | **Source** | **References** |
| --- | --- | --- | --- |
| **JIM5** | HG, low DM, partially methyl-esterified or de-esterified | Rat |  |
| **JIM7** | HG, high DM | Rat |  |
| **LM7** | HG, partially methyl-esterified (non-blockwise de-esterification) | Rat |  |
| **LM18** | HG, low DM, partially methyl-esterified or de-esterified | Rat |  |
| **LM19** | HG, low DM, preferably de-esterified (more selective than JIM5) | Rat |  |
| **LM20** | HG, high DM (more selective than JIM7) | Rat |  |
| **LM8** | Xylogalacturonan | Rat |  |
| **CCRC-M13** | RG-I | Mouse | Pattathil *et al.*, 2010 |
| **INRA-RU1** | RG-I backbone (min. 6 disaccharide repeats) | Mouse |  |
| **INRA-RU2** | RG-I backbone (min. 2 disaccharide repeats) | Mouse |  |
| **LM5** | (1→4)-β-D-galactan (min. 3 galactose units at non-reducing end) | Rat |  |
| **LM6** | (1→5)-α-L-arabinan / AGP epitopes | Rat |  |
| **LM6-M (BR12)** | (1→5)-α-L-arabinan (higher affinity than LM6) | Rat | Cornuault *et al.*, 2017 |
| **LM13** | Specific subset of unbranched (1→5)-α-L-arabinan (arabinanase sensitive) | Rat |  |
| **LM16** | Processed (1→5)-α-L-arabinan (galactosidase sensitive) | Rat |  |
| **LM26** | (1→4)-β-D-galactan substituted with (1→6)-β-D-galactosyl (branched galactan) | Rat |  |

**Table S2.** List of primary mAbs targeting hemicelluloses. Adapted from Rydahl *et al.* (2018) and references therein, unless otherwise indicated.

| **mAb** | **Antigen / Epitope** | **Source** | **References** |
| --- | --- | --- | --- |
| **BS-400-2** | (1→3)-β-D-glucan [Callose and laminarin] | Mouse |  |
| **BS-400-3** | (1→3),(1→4)-β-D-glucan [Mixed-linkage glucan, MLG] | Mouse |  |
| **CCRC-M1** | α-L-fucosylated xyloglucan / RG-I | Mouse |  |
| **CCRC-M39** | α-L-fucosylated xyloglucan / RG-I | Mouse | Pattathil *et al.*, 2010 |
| **CCRC-M58** | Xyloglucan (XLLG motif) | Mouse | Pattathil *et al.*, 2010 |
| **LM15** | Xyloglucan (XXXG motif), non-fucosylated | Rat |  |
| **LM24** | Xyloglucan (XLLG motif) | Rat |  |
| **LM25** | Xyloglucan (XLLG, XXLG, XXXG motifs) | Rat |  |
| **INRA-AX1** | Backbone of xylans | Mouse |  |
| **INRA-UX1** | Alkali-treated glucuronoxylan, GlcA (or its 4-*O*-methyl ether) substituents | Mouse | Koutaniemi *et al.*, 2012 |
| **LM10** | (1→4)-β-D-xylan | Rat |  |
| **LM23** | Non-acetylated xylosyl residues, pectic xylogalacturonan and xylan | Rat |  |
| **LM27** | Grass glucuronoarabinoxylan (GAX) | Rat |  |
| **BS-400-4** | (1→4)-β-D-(galacto)mannan | Mouse |  |
| **CCRC-M170** | Acetylated glucomannan | Mouse |  |
| **LM21** | (1→4)-β-D-(galacto)(gluco)mannan; DP2 to DP5 | Rat |  |
| **LM22** | (1→4)-β-D-(gluco)mannan; DP2 to DP5 | Rat |  |

**Table S3.** List of primary mAbs targeting glycoproteins and cell wall phenolics. Adapted from Rydahl *et al.* (2018) and references therein, unless otherwise indicated.

| **mAb** | **Antigen / Epitope** | **Source** | **References** |
| --- | --- | --- | --- |
| **JIM11** | Extensin (periodate sensitive) | Rat |  |
| **JIM12** | Extensin (proteinase sensitive) | Rat |  |
| **JIM19** | Extensin (periodate sensitive) | Rat |  |
| **JIM20** | Extensin (periodate sensitive) | Rat |  |
| **LM1** | Extensin / Hydroxyproline-rich glycoproteins (HRGPs) | Rat |  |
| **LM3** | Extensin | Rat | Feng *et al.*, 2014 |
| **JIM4** | AGP (β-D-GlcA-(1→3)-α-D-GalA-(1→2)-α-D-Rha competes for binding) | Rat |  |
| **JIM8** | AGP, carbohydrate portion | Rat | Pennell *et al.*, 1991 |
| **JIM13** | AGP (β-D-GlcA-(1→3)-α-D-GalA-(1→2)-α-D-Rha competes for binding) | Rat |  |
| **JIM16** | AGP, (1→3)-β-D-galactan chain with single (1→6)-β-D-linked Gal residue | Rat |  |
| **LM2** | AGP, (1→6)-β-D-galactan chain with terminally attached GlcA | Rat |  |
| **LM14** | AGP / Arabinogalactan | Rat |  |
| **LM30** | AGP (arabinofuranosidase sensitive) | Rat | Wilkinson *et al.*, 2017 |
| **MAC207** | AGP (β-D-GlcA-(1→3)-α-D-GalA-(1→2)-α-D-Rha competes for binding) | Rat |  |
| **LM9** | Feruloylated (1→4)-β-D-galactan | Rat |  |
| **LM12** | Feruloylate/ferulic acid on any polymer and heteroxylan | Rat |  |
